# Differentiation of Human Colon Tissue In Culture: Effects of calcium on trans-epithelial electrical resistance and tissue cohesive properties

**DOI:** 10.1101/744979

**Authors:** Shannon D McClintock, Michael K Dame, Aliah Richter, Durga Attili, Sabrina S Silvestri, Maliha M Berner, Margaret S Bohm, Kateryna Karpoff, Caroline McCarthy, Jason R Spence, James Varani, Muhammad N Aslam

**Author notes:** Corresponding author (Muhammad N Aslam).

## Abstract

**Background and aims:** Human colonoid cultures maintained under low-calcium (0.25 mM) conditions undergo differentiation spontaneously and, concomitantly, express a high level of tight junction proteins, but not desmosomal proteins. When calcium is included to a final concentration of 1.5 – 3.0 mM (provided either as a single agent or as a combination of calcium and minerals), there is little change in tight junction proteins but a strong up-regulation of desmosomal proteins and an increase in desmosome formation. The aim of this study was to assess functional consequences of the differences in calcium-mediated barrier protein expression.

**Methods:** Human colonoid-derived epithelial monolayers were interrogated in transwell culture under low- or high-calcium conditions. Ion permeability and monolayer integrity were assessed by measuring trans-epithelial electrical resistance (TEER) across the confluent monolayer. Colonoid cohesiveness was assessed in parallel.

**Results:** TEER values were high in the low-calcium environment and increased only modestly in response to calcium. In contrast, colonoid cohesiveness increased substantially with calcium supplementation. In both assays, the response to Aquamin was greater than the response to calcium alone. However, differences between interventions were small and only compared to the 0.25 mM calcium were they statistically significant. Consistent with these findings, occludin expression (a measure of tight junctions) was high at 0.25 mM calcium and did not increase with supplementation. Cadherin-17 and desmoglein-2 were weakly-expressed under low calcium conditions but increased with intervention.

**Conclusions:** These findings indicate that low ambient calcium levels are sufficient to support formation of a permeability barrier in the colonic epithelium. Higher calcium levels are necessary to promote tissue cohesion and enhance barrier function. These findings may help explain how an adequate daily intake of calcium contributes to colonic health by improving barrier function, even though there is little change in colonic histological features over a wide range of calcium intake levels.

## Introduction

An intact colonic barrier is necessary for gastrointestinal health (1–6). The intact barrier prevents permeation of toxins, soluble antigens, lipopolysaccharides and other inflammatory initiators across the intestinal wall, while inhibiting infiltration of bacteria and other particulate matter into the interstitium. Intestinal barrier defects are seen in conjunction with inflammation in the gastrointestinal tract. While inflammation itself can give rise to these defects, pre-existing weaknesses in the gastrointestinal barrier could predispose the tissue to inflammation.

Multiple cell surface structures contribute to an effective colonic barrier. Most attention has been focused on tight junctions as these structures are found at the apical surface of the mucosal epithelium and form a “seal” between adjacent cells (7,8). Along the lateral surface (i.e., beneath the apical surface) are desmosomes (9,10), which provide anchoring sites for intermediate filaments and are necessary for cohesive strength. In addition to these two cell-cell adhesional complexes are the adherens junctions (11–13). These junctional complexes are comprised of cadherin family members and mediate homotypic cell-cell attachment. Their formation between adjacent epithelial cells is a rapid event and thought to be necessary for the formation and organization of other junctional complexes. They may also contribute to cohesive strength directly or indirectly. In addition to cell-cell adhesion molecules, components of the basement membrane and moieties that mediate interactions between cells and the extracellular matrix also contribute to tissue barrier properties – especially in regard to cell trafficking and macromolecule permeability (14,15).

Calcium is critical to cell-cell and cell-matrix adhesion. Most of the molecules that mediate cellular adhesive functions depend on precise levels of calcium for homotypic and heterotypic interaction (16–18). Equally important, calcium is critical to epithelial cell differentiation (19) and many of the proteins that make up the various adhesion complexes are up-regulated as part of the differentiation response.

In recent studies, we established colonoid cultures from histologically-normal human colon tissue as well as from several large adenomas obtained at endoscopy (20,21). Once established, the colonoid cultures were maintained in a low-calcium (0.15-0.25 mM) environment or exposed to levels of extracellular calcium (1.5 – 3.0 mM) that are known to foster epithelial cell differentiation. Calcium was provided either alone or as part of Aquamin, a multi-mineral natural product, that also contains a high level of magnesium and detectable amounts of up to 72 additional trace elements in addition to calcium (22). To summarize findings from these studies, tumor-derived colonoids maintained an undifferentiated phenotype in low-calcium conditions but underwent a robust differentiation response with calcium supplementation. In contrast, normal tissue colonoids underwent differentiation in the low-calcium environment and additional calcium had only incremental effects on morphological features and differentiation marker (CK20) expression. As part of our study with histologically-normal colonoids, a combination of proteomic analysis and immunohistology was used to assess the effects of intervention on cellular adhesion molecule expression. In parallel, transmission electron microscopy was used to visualize adhesion structures. Treatment with either calcium alone or Aquamin had only modest effects on tight junction protein expression but substantially increased desmosomal proteins and cadherin family members. Basement membrane proteins were also up-regulated. Adhesion structures, especially desmosomes, were prominent in the treated colonoids. Calcium from either source was comparably effective in stimulating cell-cell adhesion proteins, but Aquamin was more effective than calcium alone at up-regulating basement membrane molecules (21).

Given the changes in adhesion protein expression, we postulated that improved barrier function and, especially, stronger tissue cohesion would be consequences of intervention. However, no data were provided to substantiate this suggestion. The present study was carried out to address this issue. Here we show that trans-epithelial electrical resistance (TEER), a measure of barrier integrity and permeability (23), was increased with calcium alone or with Aquamin supplementation but, consistent with the modest change in occludin expression, the increase was also slight. In contrast, tissue cohesion was increased substantially in response to the same interventions. This was accompanied by a strong up-regulation of desmoglein-2 and cadherin-17.

## Materials and Methods

### Calcium sources

Calcium Chloride was obtained as a 0.5 M solution (PromoCell GmbH, Heidelberg, Germany). Aquamin, a calcium-rich and magnesium-rich multi-mineral product obtained from the skeletal remains of the red marine algae, *Lithothamnion sp* (22) was provided by Marigot Ltd (Cork, Ireland) as a powder and has been used in previous studies (20,21,24,25). Aquamin contains calcium and magnesium in a ratio of approximately 12:1, along with measurable levels of 72 other trace minerals (essentially all of the minerals algae fronds accumulate from the ocean water). Mineral composition was established via an independent laboratory (Advanced Laboratories; Salt Lake City, Utah) using Inductively Coupled Plasma - Optical Emission Spectrometry (*ICP-OES*). Aquamin is sold as a dietary supplement (GRAS 000028) and is used in various products for human consumption in Europe, Asia, Australia, and North America. A single batch of Aquamin^®^ Soluble was used for this study. The complete minerals composition can be found in S1 table.

### Colonoid culture

Histologically normal colon tissue in colonoid culture was available from four subjects of our previous studies (20, 21). The collection and use of human colonic tissue was approved by the Institutional Review Board (IRBMED) at the University of Michigan. This study was conducted according to the principles stated in the Declaration of Helsinki. All subjects provided written informed consent prior to flexible sigmoidoscopy. For the present study, cryopreserved samples were re-established in culture and expanded over a 3-4 week period in growth medium as described by Dame et al. (26). Growth medium consisted of 50% L-WRN (27) conditioned Advanced DMEM/F12 (Invitrogen) providing a source of recombinant Wnt3a, R-spondin-3, and Noggin and was supplemented with 1X N2 (Invitrogen), 1X B-27 without vitamin A (Invitrogen), 1 mM N-Acetyl-L-cysteine, 10 mM HEPES (Invitrogen), 2 mM Glutamax (Invitrogen), 10 μM Y27632 (Tocris), 500 nM A83-01 (Tocris), 10 μM SB202190 (Sigma), 100 μg/mL Primocin (InvivoGen), and 100 ng/mL EGF (R&D). In addition, 2.5 μM CHIR99021 (Tocris) was used at passaging.

### TEER assay

One day prior to cultivation on transwell membranes, the colonoids were treated with a 1:1 mix of growth medium and IntestiCult-Human culture medium (StemCell Technologies). Colonoids were then dissociated into small cell aggregates (less than 40μm in size) with 0.05% Trypsin-EDTA (Invitrogen) containing 10 μM Y27632 (3.75 minutes at 37°C) and plated onto collagen IV (Sigma)-coated transwells (0.4 μm pore size, 0.33cm^2^, PET, Costar) at 200,000 individual aggregates per well (28,29). Cells were seeded for attachment and initial growth in growth medium. After 24 hours, the growth medium was replaced with KGM-Gold (Lonza), a serum-free, calcium-free medium designed for epithelial cell growth. KGM-Gold supplemented with calcium (calcium chloride or Aquamin) to bring a final concentration to 0.25, 1.5 or 3.0 mM was used for comparison whereas KGM-Gold containing 0.25 mM calcium chloride was used as control medium. As an indicator of monolayer formation, a complete differentiation medium consisting of the basal medium Advanced DMEM/ F12 media (1.05 mM calcium) and lacking the stem cell support components Wnt3a, R-spondin-3, CHIR99021, A83-01, and SB202190 was used (29). Both the treatment medium and the control differentiation medium were supplemented with 10 nM Gastrin (Sigma), 50 ng/mL Noggin (R&D), 50 ng/mL EGF, and 2.5 μM Y27632. After 2 days in differentiation medium or treatment medium, Y27632 was also removed. Media were refreshed every two days during the assay period. TEER values were determined on days-2 and −5 with an Epithelial Volt ohm meter 2 (EVOM2) and STX2 series chopstick electrodes (World Precision Instruments).

### Confocal fluorescence microscopy

At the completion of the TEER assay, membranes were prepared for confocal fluorescence microscopy. The membranes were fixed for 15 minutes at −20°C in methanol. They were then washed three times in PBS before blocking in 3% BSA (A8806; Sigma) in PBS for 1 hour. Following this, membranes were stained with antibodies to occludin (331594; Invitrogen; 1:400), desmoglein-2 (53-9159-80; eBioscience; 1:200), and cadherin-17 (NBP2-12065AF488; Novus Biologicals; 1:200) for 1 hour in 1% BSA in PBS. Stained membranes were rinsed 3 times (5 minutes each) in PBS, stained with DAPI for 5 minutes to identify nuclei and washed an additional 3 times with PBS. Finally, the membranes were gently cut from the transwell insert and mounted apical side up on Superfrost Plus glass slides (Fisher Scientific, Pittsburgh, PA) with Prolong Gold (P36930; Life Technologies Molecular Probes). The stained specimens were visualized and imaged with a Leica Inverted SP5X Confocal Microscope System (University of Michigan Medical School Biomedical Research Core Facility).

### Tissue cohesion assay

After establishment and culture expansion, colonoids were incubated for a two-week period in L-WRN conditioned Advanced DMEM diluted 1:4 with KGM Gold. The final serum concentration in the medium was 2.5% and the calcium concentration was 0.25 mM. Since the calcium is present as a component of the L-WRN conditioned medium, Aquamin could not be used in this assay at 0.25mM calcium concentration. In parallel, colonoids were incubated in the same medium supplemented with calcium (alone or as part of Aquamin) to a final concentration of 1.5 or 3.0 mM calcium. At the end of the two-week incubation period, phase-contrast microscopy (Hoffman Modulation Contrast - Olympus IX70 with a DP71 digital camera) was used to capture images in order to measure the size of multiple individual colonoids (80-140 per condition). Colonoids were then separated from the Matrigel, fragmented with mechanical force alone by pipetting the entire pellet 30x through an uncut 200 microliter pipet tip, washed 3x in PBS, and then re-cultured in fresh Matrigel. One day after establishment, multiple colonoids were again examined under phase-contrast microscopy and sized. Phase contrast images were analyzed using area measurements in Adobe Photoshop (CC version 19.1.5). Average colonoid size reduction (fold-change before and after fragmentation) was determined by dividing the average pre-fragmentation surface area by the average post-fragmentation area.

### Statistical analysis

Means and standard deviations were obtained for discrete values in both assays. Data generated in this way were analyzed by ANOVA followed by paired t-test (two-tailed) for comparison using GraphPad Prism version 8.

## Results

### Barrier integrity (measured by TEER)

Fig 1 demonstrates TEER values obtained with colonoid cells at day-2 and day-5 from 4 independent assays. At the early time-point, resistance values were low under all experimental conditions, though high in complete differentiation medium (providing maximum response). By day-5, substantial electrical resistance across the cell layer was seen under all experimental conditions. Values ranged between 51% of the maximum value with 0.25% calcium and 71% of the maximum value with Aquamin providing 3.0 mM calcium. When Aquamin-treated colonoid cells were compared to cells treated with calcium alone at comparable calcium levels, Aquamin values were 16% higher than calcium alone at 0.25 mM (p=0.018), 6% higher at 1.5 mM, and <1% higher at 3.0 mM. The TEER values from all conditions were significantly higher as compared to the control (0.25 mM Calcium).

**Fig 1.**
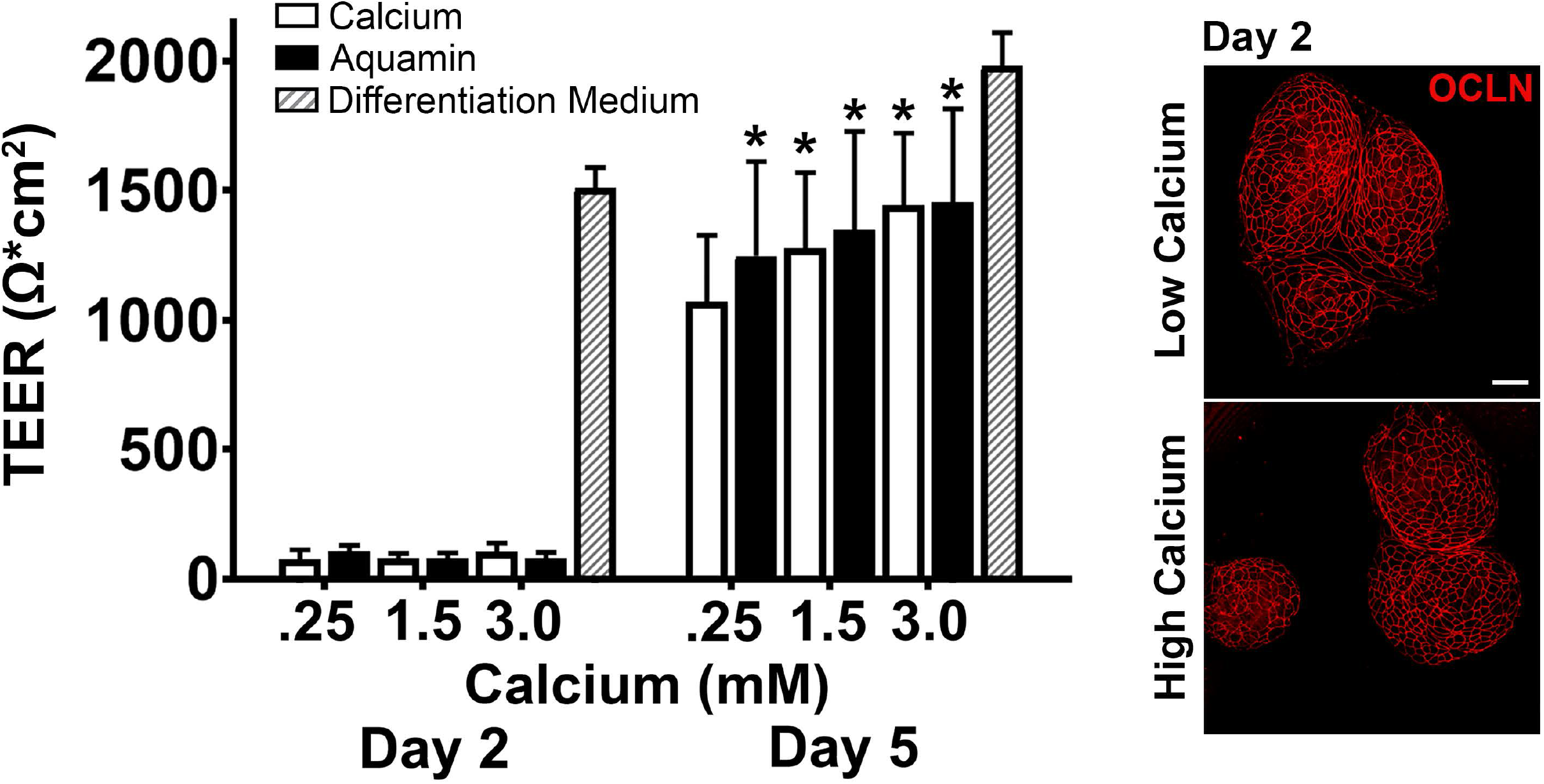
Trans-epithelial electrical resistance (TEER) values. Values shown are means and standard deviations based on four separate experiments with 3 or 4 samples (individual membranes) per data point in each experiment). Membrane to membrane variability was routinely less than 5%. Maximal TEER values represent data obtained in complete differentiation medium run in parallel with each experiment. Data were analyzed for statistical differences using ANOVA followed by paired-group comparisons. * indicate statistical significance from calcium at 0.25 mM by p<0.05. Inset: Confocal fluorescent microscopic (max projected) images of membranes stained after the day-2 reading with antibody to occludin. Scale bar = 10 μm.

At the completion of the electrical resistance measurements on day-2 and day-5, membranes were fixed and stained in the transwell inserts. The membrane-bound colonoid cells were stained with antibodies to occludin and desmoglein-2 (single membrane) or with antibodies to cadherin-17 (separate membrane) and viewed by confocal fluorescence microscopy. The inset to Fig 1 shows occludin staining at day-2 as a way to identify areas of the membrane to which cell clusters had attached and areas of the membrane that were devoid of cells. It is evident from the two images that large portions of the membrane (black areas) were devoid of cells. Thus, the lack of electrical resistance at day-2 reflects the fact that a monolayer of cells had not yet formed from the attached cell clusters (red stained). It is also evident from the two images that occludin staining was strong at 0.25 mM calcium on day-2 and not perceptibly different with calcium levels as high as 3.0 mM.

Day-5 staining results with all three antibodies are shown in Fig 2. Panels in Fig 2 a-f present findings with occludin. By day-5, an intact staining boundary could be seen between virtually all cells under all conditions. Staining intensity did not vary with the condition. The staining pattern at day-5 was indistinguishable from that at day-2. The staining pattern with desmogelin-2 was very different (panels in Fig 2 g-l). There was detectable staining with both calcium alone and Aquamin at 0.25 mM. In both cases, however, staining was patchy and punctate; no intact staining boundary was detectable between cells. At 1.5 mM, strong staining was observed throughout the monolayer and an intact boundary between adjacent cells was evident. Though differences between the two interventions were generally minute, a few areas of the membrane from the calcium-alone treatment continued to demonstrate patchy and punctate staining. At 3.0 mM, strong and uniform staining was seen throughout the cell layer with both interventions.

**Fig 2.**
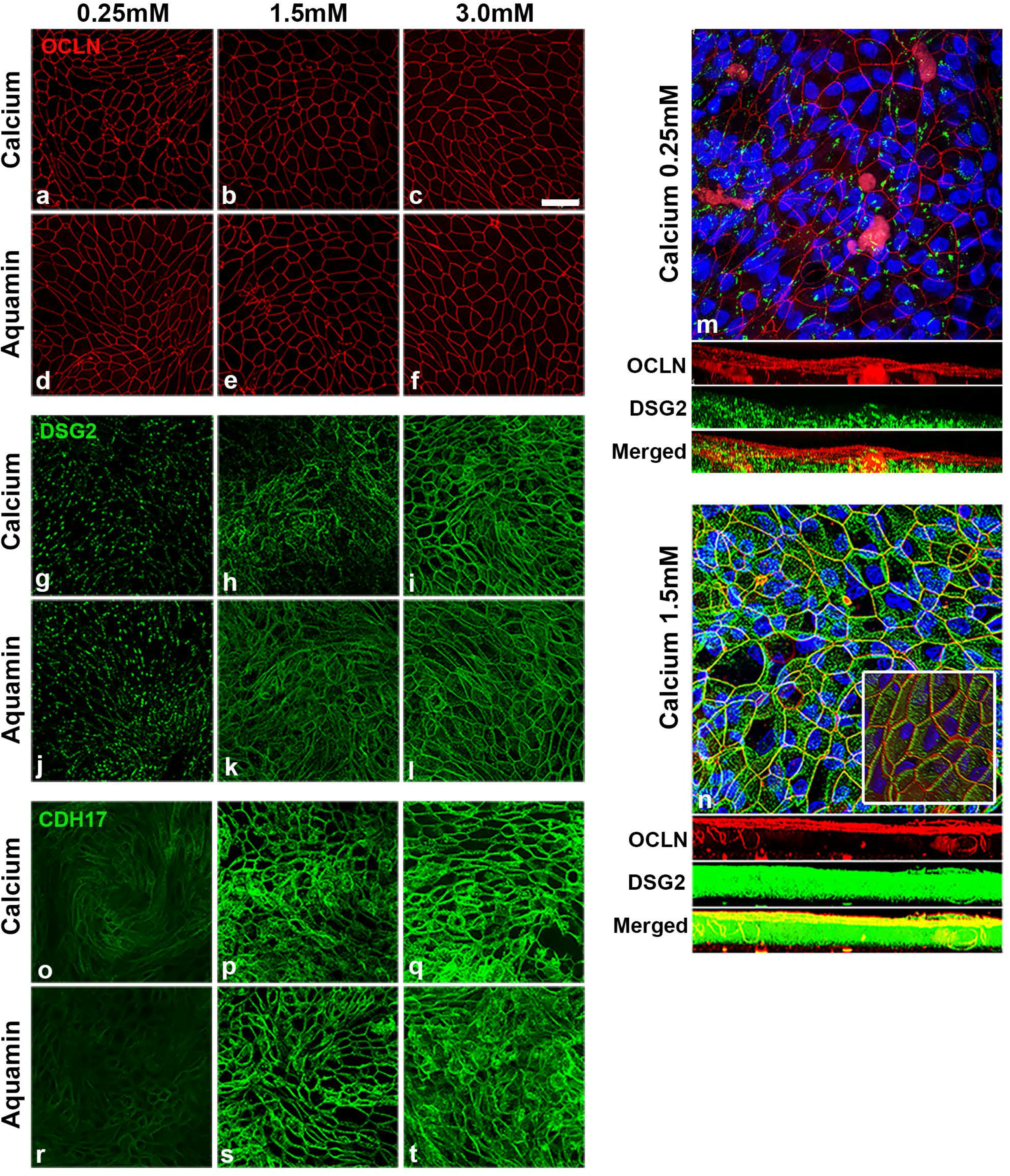
Confocal fluorescent microscopic images stained after the day-5 reading. **a-f:** Occludin (max projected); **g-l:** Desmoglein-2 (max projected). **m-n:** Occludin and desmoglein-2 (z-stack composites made up of approximately 50 planes per image. Nuclear staining is indicated by blue color (DAPI). The three bars below the main images represent horizontal views of staining through the entire membrane at each z-plane. Occludin is red, desmoglein-2 is green and the bright yellow color in the horizontal view of the 1.5 mM calcium image represents a composite of the two proteins. Inset: The inset in the 1.5 mM calcium image is a view of the entire z-stack viewed from a 45° angle to the cell surface in order to provide a 3-dimensional rendering. **o-t:** Cadherin-17 (max projected). Scale bar = 10 μm.

Panels in Fig 2 m and n show staining at 0.25 mM calcium and at 1.5 mM calcium with antibodies to both occludin and desmoglein-2. For this image, the entire “z-plane” stack is shown. The three bars shown below each image represent z-planes (approximately 50 individual planes) viewed horizontally through the cell layer from top to bottom. Under both low- and high-calcium conditions, virtually all of the red staining (occludin) is in the upper-most planes. There is little difference between the two calcium levels. With desmoglein-2 (green staining), calcium concentration-related differences are evident. At 0.25 mM calcium, most of the staining is in the lower planes while at 1.5 mM calcium, staining is distributed through the entire cell layer. The lowest bar presents a merger of the two colors. It appears from the bright yellow color at 1.5 mM calcium that both proteins are present at the cell surface. Desmoglein-2 has been shown to cycle through the apical surface during barrier formation (30), and it is possible that the two proteins are, in fact, co-localized at the apical surface. Alternatively, the monolayer is not a geometrically-flat surface and some of the upper-most z-planes may include both apical surface and sub-surface views from multiple individual cells.

The insert helps to resolve this issue. It is a high-power view of the same stack of z-planes viewed at a 45-degree angle to provide a 3-dimensional representation. With occludin (red stain), the cell-cell boundary can be seen at the apical surface of the cells over the entire cell layer. Desmogelin-2 (green stain) is seen at the surface but immediately below occludin, and extends downward along the lateral boundary between adjacent cells.

Using a separate transwell membrane, colonoid cells (day-5) were stained with antibody to cadherin-17. Results shown in Fig 2 o-t panels indicate low-level cadherin-17 expression in 0.25 mM calcium and a dramatic up-regulation with calcium alone (1.5 or 3.0 mM) or with Aquamin providing the same amount of calcium. Of interest, and unlike what was observed with desmoglein-2 at 0.25 mM calcium, cadherin-17 expression in the low-calcium environment was not patchy or punctate. Rather, the cell border was well-defined, even though only faintly visible.

### Colonoid cohesion

Fig 3 presents results from colonoid cohesion studies. For these studies, colonoids were maintained for a 14-day period in culture medium (LWRN/KGM Gold) containing either 0.25 mM calcium (control) or supplemented with additional calcium. At the end of the incubation period, surface area measurements were made on multiple individual colonoids as described in the Materials and Methods Section. Following this, subculture was done and individual colonoids were “sized” again. With colonoids maintained in culture medium containing 0.25 mM calcium, there was an approximately 9-fold reduction in colonoid size during subculture (i.e., average post-split size of individual colonoids compared to average pre-split size). In comparison, colonoids incubated in culture medium containing 1.5 or 3.0 mM calcium demonstrated only 5.2-fold and 4.2-fold reductions in size. With Aquamin, pre-split versus post-split differences in colonoid size were even smaller (i.e., 3.7-fold and 3.5-fold reductions).

**Fig 3.**
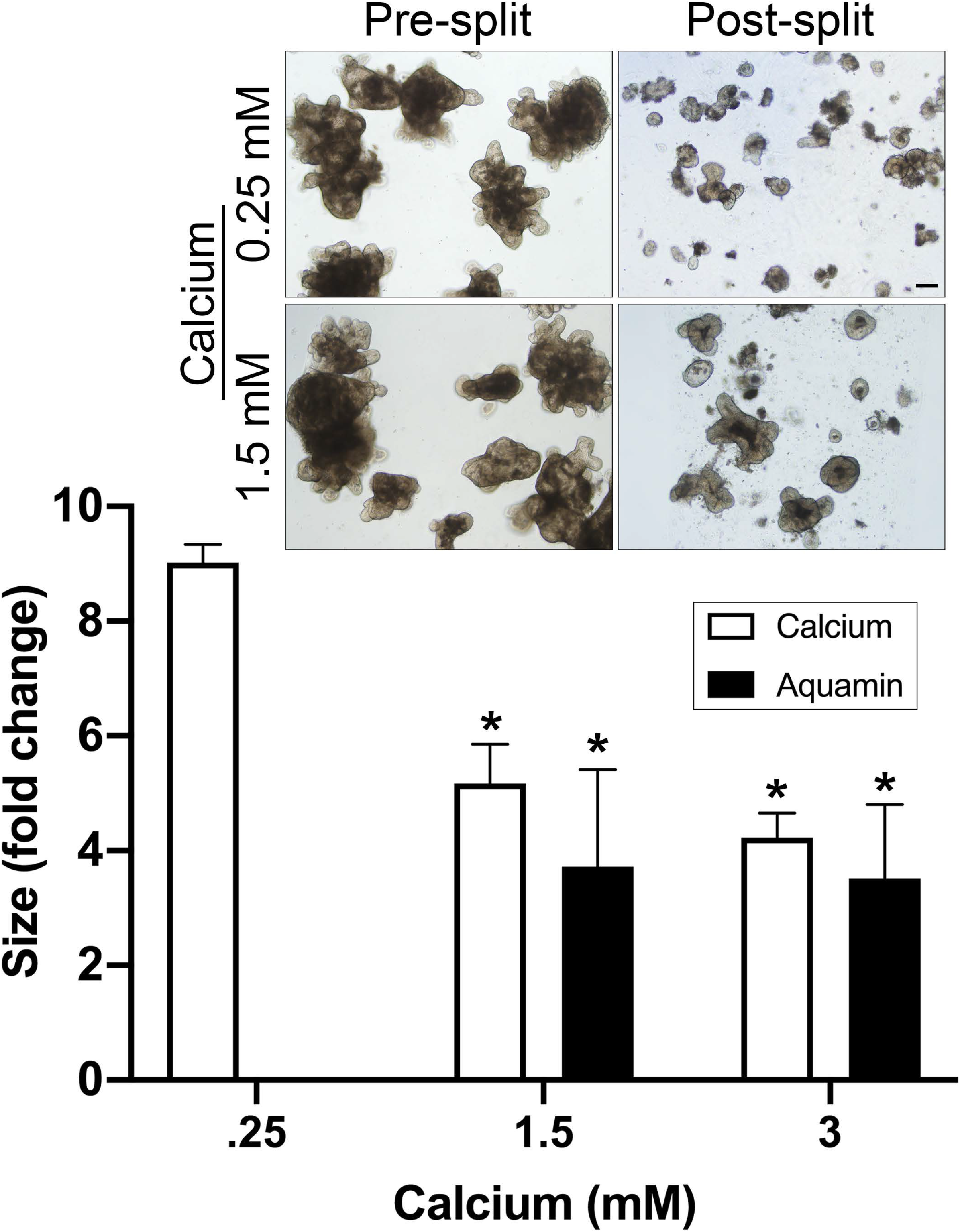
Colonoid cohesion. Values shown are change in the surface area means and standard deviations of individual colonoids based on three separate experiments (representing colonoids from three different subjects) with a minimum of 80 colonoids assessed per treatment group in both pre- and post-split cultures. Data were compared for statistical differences using ANOVA followed by paired-group comparisons. * indicates difference from 0.25 mM calcium alone at p<0.05. Inset: Representative examples of pre-split and post-split colonoids. Scale bar = 200 μm.

## Discussion

Calcium is the quintessential promoter of epithelial cell differentiation (19). Calcium levels above approximately 1.0 mM are required for differentiation to occur with epithelial cells in monolayer culture. It was surprising, therefore, when our recent studies (20,21) demonstrated that differentiation (indicated by colonoid gross and histological appearance and by differentiation marker [CK20] expression) was seen in normal human colonoids maintained under low-calcium (0.25 mM) conditions. In these colonoids, the tight junctional protein, occludin, was detectable at the cell surface by immunostaining and tight junctions were evident by electron microscopy. In parallel, proteomic analysis showed that a number of tight junction proteins (claudin-3 and −4, as well as occludin) were only minimally up-regulated with calcium supplementation as compared to control (0.25 mM calcium). In contrast, however, a number of other cell surface and extracellular matrix proteins were minimally expressed under low-calcium conditions, but substantially up-regulated with calcium supplementation. Among these were desmosomal proteins (desmoglein-2, desmocollin-2 and desmoplakin), cadherin family members (cadherin-17, protocadherin-1, cadherin-related family members-2 and −5), and proteins found at the apical surface of the colonic epithelium (mucins and CEACAMs). Additionally, several basement membrane components (laminin A and B chains, nidogen and heparin sulfate proteoglycan) were also strongly up-regulated in response to calcium supplementation. The significance of these calcium concentration-related differences in protein expression to functional behavior was not addressed. In the present study, we used trans-epithelial cell electrical resistance across colonoid-derived epithelial cells in 2-dimensional transwell membranes as a measure of tissue integrity and permeability. An assay for tissue cohesion was employed in intact 3-dimensional colonoids. Functional responses were compared with protein expression data.

Electrical resistance across the polarized gastrointestinal epithelial layer correlates with existence of a barrier to small molecule passage between adjacent epithelial cells (2–4); it is generally accepted that tight junctions, primarily, mediate this property (23). Consistent with occludin expression data - both in the intact colonoids (21) and cells derived from the colonoids (this report) - trans-epithelial electrical resistance was substantial under low-calcium conditions and was increased only modestly in response to calcium supplementation. Thus, it appears that the low calcium level needed to initiate differentiation in colonoid culture is also sufficient to support permeability barrier formation. Formation of tight junctions between adjacent epithelial cells follows cell-cell adhesion and formation of adherens junctions. Cadherin-1 (E-cadherin) is primarily responsible for this initial adhesive interaction (31,32). This is of interest because although several cadherin family members were induced with calcium supplementation, cadherin 1 was not one of them. Rather, cadherin-1 was highly expressed in the low-calcium environment (21,33) and not further induced with calcium supplementation.

While trans-epithelial electrical resistance was generated under low-calcium conditions, the same low calcium level did not support strong tissue cohesion. Rather, a substantial increase in cohesion was observed with either calcium alone or Aquamin at 1.5 and 3.0 mM as compared to the low-calcium control. Increased cohesion correlated with up-regulation of desmosomal proteins (citation 21 and this report) and with an increase in actual desmosomes seen at the ultrastructural level (21). Of interest, increased cadherin-17 expression occurred concomitant with elevated expression of desmoglein-2 While cadherin-17 does not participate in the formation of adherens junctions, this cadherin is highly responsive to small changes in calcium concentration and does contribute to homotypic and heterotypic cell adhesion in the intestinal epithelium (34). Cadherin-17 up-regulation with calcium supplementation in parallel with desmosomal protein up-regulation could help organize desmosomal proteins into the structures that are responsible for strong cell-cell cohesion.

While the relationship between desmosome formation and tissue cohesion is clear, the role these structures play in the regulation of small-molecule permeability is less so. Perhaps the increase in electrical resistance seen with calcium supplementation is a reflection of a direct contribution of desmosomes to the permeability barrier. A recent study demonstrated a signaling role for desmogelin-2 in tight junction formation during barrier repair (30). Alternatively, desmosome contribution may be indirect. It is difficult to envision precise control of permeability in the absence of strong tissue cohesion – especially in a mechanically-active tissue such as the colon. Additional studies will be needed to address this complex issue.

Along with molecules that directly affect cell-cell adhesion, our recent study (21) also demonstrated up-regulation of several other moieties that may contribute to overall barrier formation in the colon. Among these are components of the basement membrane (laminin A and B chains, nidogen and heparan sulfate proteoglycan). In a simple epithelium such as the colon, there is only one layer of epithelial cells, and every cell resides on the basement membrane. If the basement membrane is not optimally constituted, or if cells cannot properly attach to the basement membrane, rapid apoptosis and sloughing of the cell layer will occur (35,36) – making more-subtle effects on barrier function moot. Furthermore, the basement membrane itself influences transit of cells and macromolecules from the colonic fluid into the interstitium (14,15).

Finally, molecules that form the carbohydrate-rich layer above the colonic epithelium (mucins and CEACAMs) were also found to be up-regulated with supplementation. These carbohydrate-rich molecules trap bacteria and other particulates, preventing them from reaching the epithelial cell surface in the first place (37,38). All of these moieties, undoubtedly, contribute to overall protective barrier in the colon.

There is a strong relationship between defective barrier function and inflammatory diseases in the gastrointestinal tract. Barrier dysfunction has been noted in both Crohn’s disease and ulcerative colitis (39–41) and is postulated as a component of irritable bowel syndrome (42) and celiac disease (43). Defective barrier function and chronic inflammation have also been seen even without overt bowel disease; e.g., as a consequence of high-fat diets (44), hyperglycemia (45), psychological stress (46) and hypoxia (47). It is well-accepted that inflammatory stimuli can provide the initial impetus for barrier breakdown (48). Once barrier defects occur, bacteria, bacterial products and other toxins / allergens can gain access to the interstitium. This promotes additional inflammation, which, in turn, leads to additional barrier breakdown. Our findings in no way contradict this understanding. They argue, however, that even in the presence of conditions that promote barrier damage in the gastrointestinal tract, having a level of calcium that supports optimal elaboration and function of key barrier proteins would be beneficial. Our data also argue that focusing only on tight junctions and small molecule permeability may be too limited to fully appreciate barrier changes in colonic disease.

In this study, calcium provided as a single agent was compared to calcium delivered as part of a calcium- and magnesium-rich multi-mineral supplement. In both the TEER and cohesion assays, responses to the multi-mineral intervention were greater than those to calcium alone. The greatest differences were seen at lower ambient calcium levels. What accounts for the enhanced response to the multi-mineral intervention is not known with certainty, but previous studies provide insight. Aquamin itself, as well as certain of the individual trace elements present in the multi-mineral product, activate the extracellular calcium-sensing receptor, a critical target in epithelial cell responses to calcium, more effectively than calcium itself (49–51). In the presence of the additional trace elements, a “left-shift” in calcium response occurs. Additional studies have demonstrated that trace elements of the lanthanide family (as present in Aquamin) have potent effects on calcium channels (52) as well as effects on calcium “pumps” (53). The behavior in one or more of these critical effectors of calcium function could interfere with calcium responses. Alternatively, rather than modulating calcium metabolism, *per se*, the major effect of the multi-mineral supplement may be directly on the adhesion process. While extracellular calcium is critical to both cell-cell and cell-matrix adhesion, other divalent cations (notably, magnesium and manganese) are also required for optimal adhesive interactions (54,55). These possible mechanisms are not mutually exclusive.

## Conclusions

These findings may help explain how an adequate daily intake of calcium contributes to health, despite their being little change in colonic histological features over a wide-range of calcium intake amounts (56–59). While having an adequate daily intake of calcium is important to health, most individuals are calcium deficient. Studies from North America, Europe and Australia have all shown that a majority of individuals do not achieve a minimal daily calcium intake (60–65). The “Western-style diet” has been thought to underlie the chronic calcium deficiency, but a recent study has shown that even where a rural agrarian diet is the norm, significant calcium-deficiency exists (66). From a public health standpoint, ensuring an adequate calcium-intake along with cofactors such as vitamin D and, perhaps, additional trace elements needed for optimal calcium uptake and function may provide a cost-effective way to improve overall health and well-being in a large segment of the population. It might be noted in this regard that in addition to the consequences of epithelial barrier dysfunction in the gastrointestinal tract, a relationship between epithelial barrier dysfunction in lung and skin has been suggested for both asthma and eczema (67,68).

A critical issue is whether and to what extent these findings in colonoid culture are reflective of what occurs *in vivo* in the intact colon. Can we, in fact, duplicate the colonoid culture findings with intervention in human subjects? We are currently in the midst of a 90-day interventional trial in healthy human adult subjects, comparing the effects of calcium alone (800 mg/day) and Aquamin (delivering 800 mg of calcium per day) to placebo for effects on the same colon biomarkers of growth, differentiation and barrier formation as assessed in colonoid culture (clintrials.gov; NCT02647671). Results of the ongoing trial will be instructive as to the relative efficacy of the multi-mineral approach compared to calcium alone as well as provide an indication of tolerability and safety. Data generated in the interventional trial (69), along with those data generated here and in our recent colonoid studies (20,21), will, ultimately, demonstrate whether multi-mineral supplementation has greater efficacy than calcium alone or whether the presence of multiple trace elements can provide efficacy at lower calcium dose.

## Acknowledgements

We thank Marigot LTD (Cork, Ireland) for providing Aquamin^®^ as a gift. We thank the Microscopy and Imaging Laboratory (MIL) for help with confocal fluorescence microscopy. We thank the Translational Tissue Modeling Laboratory (TTML) for help with colonoid propagation and help with the TEER assay. The TTML is a University of Michigan funded initiative (Center for Gastrointestinal Research, Office of the Dean, Comprehensive Cancer Center, Departments of Pathology, Pharmacology, and Internal Medicine) with support by the Endowment for Basic Sciences.

---

**S1 Table.** Aquamin^®^ Mineral Analysis.

| Element | µg/g | Element | µg/g | Element | µg/g |
| --- | --- | --- | --- | --- | --- |
| Aluminum | 21.6 | Hafnium | 0.038 | Rubidium | 0.031 |
| Antimony | 0.69 | Holmium | 0.010 | Ruthenium | 0.137 |
| Arsenic | 0.239 | Indium | <0.001 | Samarium | 0.037 |
| Barium | 1.76 | Iodine | 1.81 | Scandium | 0.469 |
| Beryllium | <0.5 | Iridium | <0.001 | Selenium | <0.5 |
| Bismuth | <0.5 | Iron | 143 | Silicon | 16.8 |
| Boron | 13.7 | Lanthanum | <0.5 | Silver | <0.5 |
| Cadmium | 0.220 | Lead | 0.084 | Sodium | 2,206 |
| Calcium | 117,000 | Lithium | <0.5 | Strontium | 882 |
| Carbon | 26,600 | Lutetium | <0.001 | Sulfur | 1,241 |
| Cerium | 0.314 | Magnesium | 10,210 | Tantalum | 0.043 |
| Cesium | 0.001 | Manganese | 25.4 | Tellurium | <0.5 |
| Chloride | 612 | Mercury | <0.001 | Terbium | 0.007 |
| Chromium | <0.5 | Molybdenum | <0.5 | Thallium | <0.5 |
| Cobalt | <0.5 | Neodymium | 0.170 | Thorium | 1.30 |
| Copper | <0.5 | Nickel | 0.75 | Thulium | 0.004 |
| Dysprosium | 0.045 | Niobium | <0.5 | Tin | 0.029 |
| Erbium | 0.033 | Osmium | <0.001 | Titanium | 11.4 |
| Europium | 0.013 | Palladium | 0.179 | Tungsten | <0.5 |
| Fluoride | 3.57 | Phosphorous | 189 | Vanadium | <0.5 |
| Gadolinium | 0.044 | Platinum | <0.001 | Ytterbium | 0.030 |
| Gallium | 0.307 | Potassium | 70.0 | Yttrium | <0.5 |
| Germanium | <0.001 | Praseodymium | 0.040 | Zinc | 6.07 |
| Gold | <0.5 | Rhenium | 0.001 | Zirconium | <0.5 |
|  |  | Rhodium | 0.061 |  |  |

Source: 2017 Test Certificate for Aquamin® Soluble, by Advanced Laboratories, Inc. (Salt Lake City), for client Marigot Limited (Ireland). The levels of individual trace elements were determined by Inductively Coupled Plasma Optical Emission Spectrometry (ICP-OES) except Carbon (determined by LECO), Chloride, Iodine (determined by Titration), and Fluoride (determined by AOAC 939.11).

